# Moyamoya Disease-Associated *RNF213* Alleles Encode Dominant Negative Alleles That Globally Impair Ubiquitylation

**DOI:** 10.1101/2020.05.24.113795

**Authors:** Abhishek Bhardwaj, Robert S. Banh, Wei Zhang, Sachdev S. Sidhu, Benjamin G. Neel

## Abstract

Single nucleotide polymorphisms (SNPs) in *RNF213*, which encodes a 591kDa protein with AAA+ ATPase and RING E3 domains, are associated with a rare, autosomal dominant cerebrovascular disorder, Moyamoya disease (MMD). MMD-associated SNPs primarily localize to the C-terminal region of RNF213, and some affect conserved residues in the RING domain. Although the autosomal dominant inheritance of MMD could most easily be explained by RNF213 gain-of-function, the type of ubiquitylation catalyzed by RNF213 and the effects of MMD-associated SNPs on its E3-ligase activity have remained unclear. We found that the RING domain of RNF213 uses the E2-conjugating enzyme UBE2D2 to catalyze predominantly K6-dependent poly-ubiquitination events comprising a mixture of typical and atypical ubiquitin linkages. MMD-associated SNPs encode proteins with decreased E3-ligase activity and the most frequent MMD allele, *RNF213*^*R4810K*^, is a dominant negative mutant that decreases ubiquitylation globally. By contrast, MMD-associated *RNF213* SNPs do not affect ATPase activity. We propose that decreased RNF213 E3-ligase activity is central to MMD pathogenesis.

## Introduction

Moyamoya disease (MMD) is a rare, autosomal dominant disorder characterized by stenotic/occlusive lesions in the circle of Willis with surrounding abnormal blood vessels in the brain (Nanba et al., 2006). These vessels have a “puff of smoke” appearance in imaging studies (hence the Japanese term, “Moyamoya”), and are believed to develop around the occlusive lesions to compensate for lack of blood flow. MMD occurs worldwide, but its prevalence and incidence are highest in East Asians (Kim, 2016). Incidence is bimodal, with peaks at ages 5-18 and 45-60, respectively (Kim, 2016). Approximately 10-40% adults and 2.8% children with MMD suffer intracranial hemorrhage, and 50-75% of patients experience a transient ischemic attack or stroke (Scott and Smith, 2009). Fifteen percent of MMD cases are familial, and these have earlier mean onset (11.8 years) compared with sporadic cases (30 years) (Nanba et al., 2006).

Single nucleotide polymorphisms (SNPs) in *RNF213* are strongly associated with MMD risk (Liu et al., 2011). On average, ∼50% of Asian (and ∼80% of Japanese) MMD families carry the *RNF213*^*R4810K*^ allele (Cecchi et al., 2014; Moteki et al., 2015). Several other rare *RNF213* variants are found in MMD patients of diverse ethnicities (Cecchi et al., 2014; Kobayashi et al., 2016; Moteki et al., 2015). Although ∼2% of Japanese individuals have *RNF213*^*R4810K*^, the prevalence of MMD is low (∼0.006%), indicating that additional genetic and/or environmental modifiers are required for pathogenesis (Ran et al., 2013).

*RNF213* encodes an ∼591kDa protein containing tandem AAA+ ATPase domains and a RING E3 domain (**Fig. 1a**). The AAA+ ATPase domains mediate RNF213 oligomerization into a homo-hexamer (Morito et al., 2014). Nucleotide (ATP/ADP) binding to the first ATPase domain stabilizes the oligomer, and destabilization occurs upon ATP hydrolysis by the second domain. Less, however, is known about the RNF213 E3 domain.

**Figure 1:**
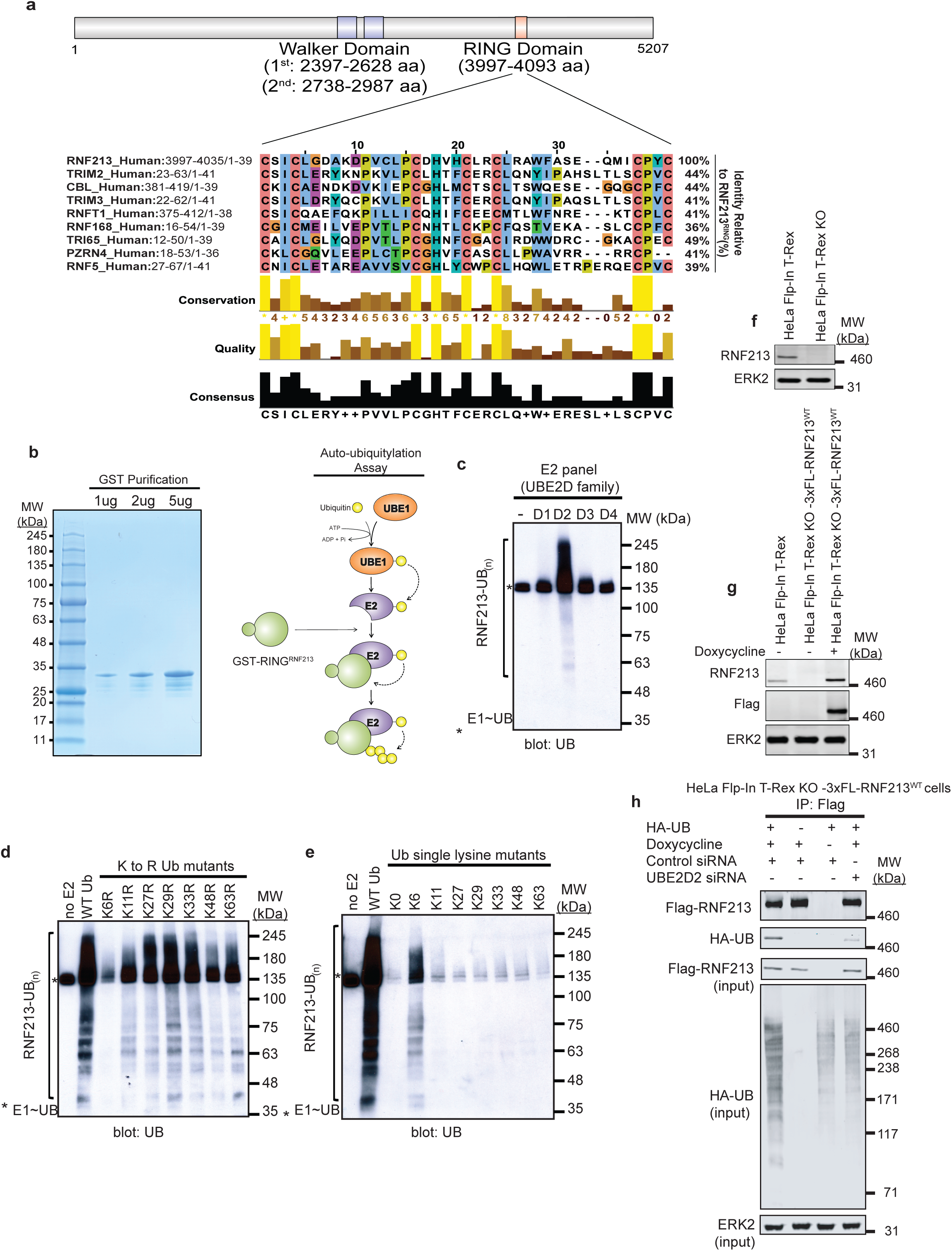
The RNF213 RING domain can catalyze K6-dependent ubiquitin linkages. **a**, Schematic of RNF213 structure, illustrating sub-domains, and multiple sequence alignment showing closest relatives of RNF213 RING. Alignment was performed by searching the RNF213 RING domain sequence against the human Uniprot database, using the default settings in Blastp. The top 8 sequences were selected a and alignment diagram was generated using Jalview. **b**, Coomassie blue stain of GST-RING^RNF213^ variants purified from *E. coli* (left panel) and scheme of *in vitro* auto-ubiquitylation assay (right panel) **c**, Auto-ubiquitylation assays using GST-RING^RNF213^ and members of the UBE2D family of E2s. **d**, GST-RNF213^RING^ has impaired auto-ubiquitylation activity in the presence of K6R ubiquitin mutant, but not other ubiquitin K>R mutants. **e**, *In vitro* auto-ubiquitylation assays were performed using GST-RING^RNF213^ and a ubiquitin mutant with all lysines mutated (K0) or single add-back (single functional lysine) mutants. Note that GST-RING^RNF213^ auto-ubiquitylation activity requires K6 on ubiquitin to form poly-ubiquitin linkages. f, Immunoblot of single cell clone shows absence of RNF213 in HeLa Flp-In T-Rex KO cells generated by CRISPR/Cas9 technology. **g**, HeLa Flp-In T-Rex KO cells from **1f** reconstituted with doxycycline-inducible 3xFL-RNF213 were induced for 36 hr, lysed, and immunoblotted for the indicated proteins. **h**, HeLa Flp-In T-Rex KO cells expressing 3x-Flag-RNF213 upon doxycycline induction were transfected with Control or *UBE2D2* siRNAs, followed by an HA-UB (HA-tagged Ubiquitin) expression construct. Cells were harvested 36 hours after transfection, with MG132 (10 µM) and Chloroquine (50 µM) added for the last 3 hours, lysed, and immunoblotted for the indicated proteins. Each immunoblot (**c-h**) is a representative image of at least three independent experiments.

RING E3 domains catalyze UB (ubiquitin) transfer from (an) E2 ubiquitin-conjugating enzyme(s) to lysine residues of E3-bound substrate(s), resulting in isopeptide bond formation (Deshaies and Joazeiro, 2009). Mono-ubiquitylation joins a single UB molecule to a substrate, although multiple sites on the same substrate can be mono-ubiquitylated (“multi-mono-ubiquitylation”). Poly-ubiquitylation occurs when UB residues are added sequentially to a specific lysine residue in a previously conjugated UB, forming a UB chain. The N-terminal methionine and seven lysine residues (K6, K11, K27, K29, K33, K48 and K63) in UB can participate in poly-ubiquitylation, leading to linear and/or branched UB chains (Yau and Rape, 2016). Some E2s appear to promote specific types of UB linkages; others are more promiscuous. Most E3 ubiquitin ligases can interact with several E2s, thereby catalyzing production of different types of UB chains. Depending on the specific UB linkage, ubiquitylation regulates diverse cellular processes, including protein degradation, endocytic trafficking, inflammation, translation and DNA repair (Yau and Rape, 2016).

Most MMD-associated SNPs map to the C-terminal region of RNF213, which includes the RING domain (Koizumi et al., 2016). Some occur in or near conserved RING residues (e.g., C3997Y, P4007R, D4013N, H4014N, R4019C). However, the type(s) of ubiquitylation catalyzed by RNF213 and whether MMD-associated SNPs affect E3 ligase activity have remained unclear.

Here, we report that RNF213 has K6-ubiquitin-dependent E3 ligase activity is mediated via the E2 UBE2D2. Furthermore, MMD-associated SNPs, including the most frequent allele, *RNF213*^*R4810K*^, encodes proteins with impaired E3-ligase activity that acts as dominant negative mutants.

## Results

### The RNF213 RING domain is dependent on K6-ubiquitin linkages

A search of the RNF213 RING against the human UniProtKB/Swiss-Prot database using Basic Local Alignment Search Tool (BLAST) (Gish and States, 1993) revealed substantial similarity to other E3 ligases (**Fig. 1a**). The strongest similarity (44% identity) was to the RING domains of TRIMs and CBL. We cloned the RNF213 RING domain (RING^RNF213^) into a bacterial expression vector to generate an N-terminal GST-tagged fusion protein, purified GST-RING^RNF213^ to homogeneity (**Fig. 1b**) and monitored auto-ubiquitylation (**Fig. 1c**). Because the E2 for RNF213-catalyzed reactions is unknown, we selected TRIM- and CBL-preferred E2s for these assays. Yeast-two hybrid screens suggest that TRIMs prefer the UBE2D and UBE2E family (Napolitano et al., 2011). CBL can interact with up to 12 E2s, but employs UBE2D family E2s to catalyze the ubiquitylation of receptor tyrosine kinases, a known physiological function of CBL (Kar et al., 2012). Moreover, a high proportion of all RING E3s interact with the UBE2D and UBE2E class of E2s (Markson et al., 2009). Therefore, we tested GST-RING^RNF213^ in concert with UBE2D family members (**Fig. 1c**). GST-RING^RNF213^ auto-ubiquitylated in the presence of UBE2D2, but not UBE2D1, UBE2D3, or UBE2D4. The products of the UBE2D2/RNF213 reaction appeared as a ladder on SDS-PAGE, suggesting that GST-RING^RNF213^ catalyzes multi-mono-ubiquitylation or poly-ubiquitylation.

We used UB mutants containing single lysine to arginine (K>R) mutations to characterize the linkages generated by GST-RING^RNF213^ (**Fig. 1d**). GST-RING^RNF213^ auto-ubiquitylation was decreased with the K11R, K27R, K33R, K48R and K63R UB mutants as substrates, but it was abrogated in the presence of UB-K6R (**Fig. 1d**). These results indicate that the RNF213 RING, using UBED2 as the E2, catalyzes non-canonical poly-ubiquitylation reactions that require an initial K6-UB linkage(s). Consistent with this notion, mutation of all UB lysines (K0) blocked RNF213 auto-ubiquitylation, whereas a ubiquitin “add-back” mutant retaining only K6, but no other single lysine add-back mutant, substantially restored activity (**Fig. 1e**). K6-alone UB did not, however, catalyzed wild type (WT) levels of auto-ubiquitylation. These findings suggest that GST-RING^RNF213^/UBED2 first generates K6 linkages, which results in enzyme activation, followed by other types of conjugation reactions.

To evaluate its physiological relevance in RNF213-mediated ubiquitination, we assessed ubiquitylation in HeLa cells with or without UBE2D2. First, we used CRISPR/Cas9 technology to generate a null (frameshift) mutant of *RNF213* in HeLa Flp-In T-Rex cells (HeLa Flp-In T-Rex KO) **Fig. 1f)**. We then used FLP-catalyzed recombination to generate HeLa Flp-In T-Rex KO-3xFL-RNF213^WT^ cells, which express wild-type 3xFlag-tagged RNF213 upon doxycycline induction **(Fig. 1g).** Finally, Hela Flp-In T-Rex KO 3xFL-RNF213^WT^ cells were co-transfected with an expression construct for HA-UB (HA-tagged Ubiquitin) and an siRNA targeting *UBE2D2* **(Fig. 1h)**. Flag-RNF213 was immunoprecipitated with anti-Flag (FL) antibody, and its ubiquitinylation was detected by immunoblotting with anti-HA antibody. In accord with our previous findings (Banh et al., 2016), RNF213 expression (in HeLa Flp-In T-Rex KO cells) increased overall ubiquitylation (compare lanes 1 and 3). Notably, *UBE2D2* knockdown, which was validated by qRT-PCR **(Fig. S1)**, reduced global ubiquitylation in RNF213-expressing HeLa cells to an extent comparable to that seen in cells lacking RNF213 expression **(Fig. 1h)**. Furthermore, consistent with our hypothesis, RNF213-ubiquitylation was reduced substantially in *UBE2D2*-knockdown cells. **(Fig 1h)**.

### Moyamoya SNPs do not affect RNF213 ATPase activity

The RNF213 AAA+ ATPase domain is required for homo-hexamer formation (Morito et al., 2014), and although R4810 is located outside of the AAA+ ATPase and RING domains, it was reported that RNF213^R4810K^ has impaired ATPase activity (Kobayashi et al., 2015). Furthermore, previous reports disagree about whether defective ATPase activity is (Kobayashi et al., 2015) or is not (Morito et al., 2014) involved in MMD pathogenesis. Both of these reports tested the ATPase activity of RNF213^R4810K^. To evaluate more generally the role of RNF213 ATPase activity in MMD pathogenesis, we quantified the *in vitro* ATPase activity of multiple MMD-associated RNF213 proteins. Using FLP-catalyzed recombination, we first generated HeLa Flp-In T-Rex KO lines expressing 3x-Flag-*RNF213*^WT^ (wild type), 3x-Flag-*RNF213*^*E2488Q,E2845Q*^ (AAA+ ATPase double mutant), 3x-Flag-*RNF213*^*I3999A*^ (RING-impaired), and four major MMD SNPs: 3x-Flag-*RNF213* ^*D4013N*^, 3x-*Flag-RNF213*^*D4014N*^, 3x-Flag-*RNF213*^*K4732T*^, and 3x-Flag-*RNF213*^*R4810K*^ **(Fig. 2a).** Each protein was purified by Flag-immunoprecipitation to near homogeneity **(Fig. 2b)** and assayed for ATPase activity **(Fig. 2c**). As expected, 3x-Flag-RNF213^E2488Q,E2845Q^ had no detectable activity. The RING mutant, 3x-Flag-RNF213^I3999A^, had ATPase activity comparable to that of 3x-Flag-RNF213^WT^, indicating that at least *in vitro*, RNF213 ATPase activity is independent of its E3-ligase activity. Similarly, the proteins encoded by MMD SNPs had wild type levels of ATPase activity, arguing that MMD is not due to defective ATPase activity, at least MMD caused by the most prevalent *RNF21*3 alleles.

**Figure 2.**
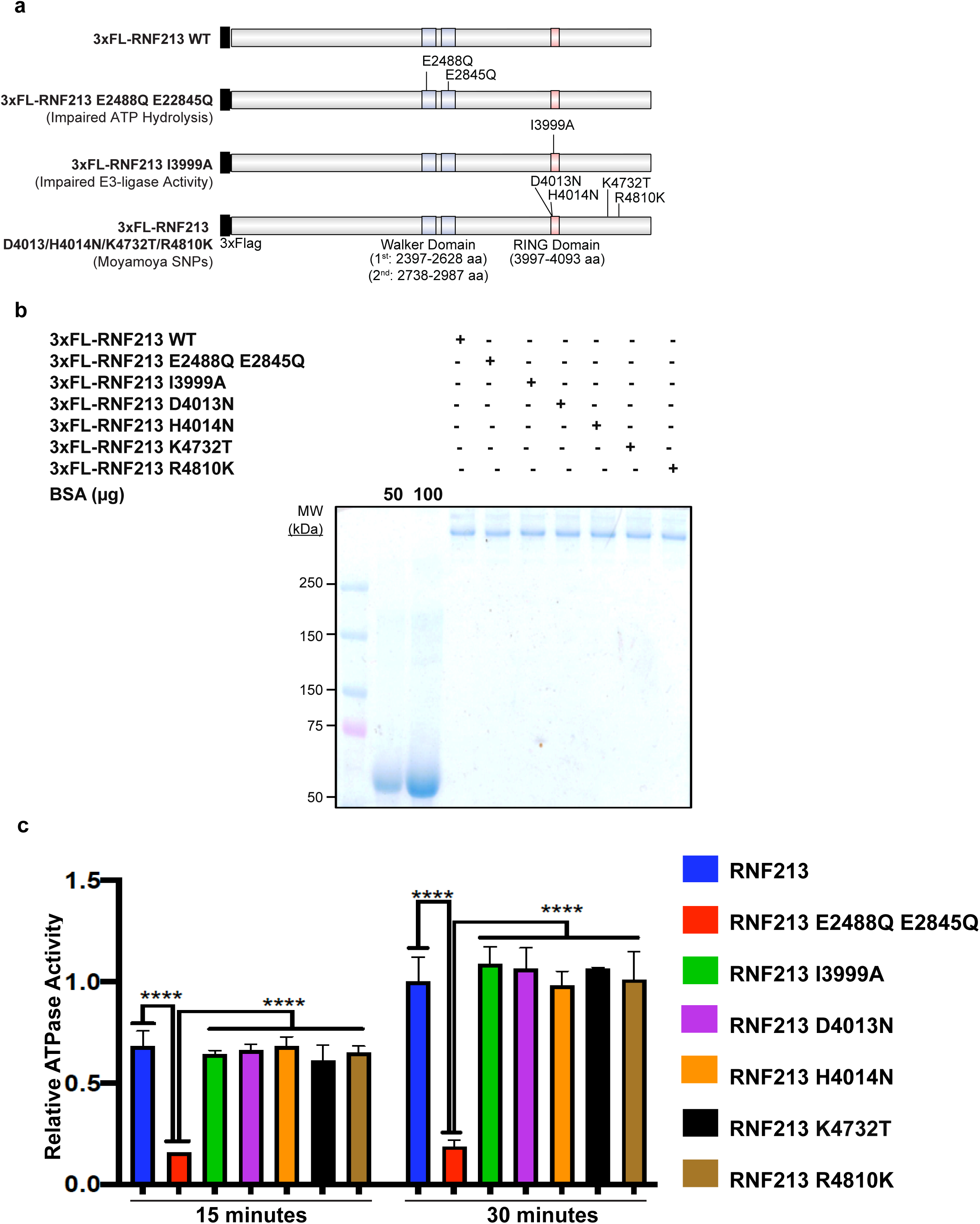
MMD SNPs do not affect AAA+ ATPase activity of RNF213. **a**, Schematic showing RNF213^WT^ and positions of AAA+ ATPase (E2488Q, E2845Q), RING (I3999A) and MMD-associated (D4013N, H4014N, K4732T, R4810K) mutants. **b**, HeLa Flp-In T-Rex KO cells were transfected with 3x Flag RNF213^WT^, 3x Flag RNF213^E2488Q, E2845Q^, 3x-Flag-RNF213^I3999A^, 3x-Flag-RNF213^D4013N^, 3x-Flag-RNF213^H4014N^, 3x-Flag-RNF213^K4732T^ or 3x-Flag-RNF213^R4810K^. Thirty-six hours post-transfection, cells were harvested and lysates were subjected to immunoprecipitation using anti-Flag antibody. Purified RNF213 variants were detected by SDS-PAGE and Coomassie blue staining. BSA was loaded to aid in quantification of RNF213. **c**, ATPase activity of purified RNF213 variants from **Fig. 2b**. Error bars represent means ± SD of three independent experiments, each with samples in triplicate. Statistical significance was evaluated by two-way ANOVA, followed by Dunnett test.

### MMD-associated SNPs encode mutants with decreased E3 activity in cells

As noted above, RNF213 is known to homo-hexamerize (Morito et al., 2014). We next asked whether MMD-associated SNPs affected RNF213 self-association. To this end, we assessed the ability of 3x-Flag-tagged (FL-)RNF213^WT^, an AAA+ ATPase-inactive mutant (E2488Q), a RING E3-defective mutant (I3999A) and the most prevalent MMD variant (R4810K) to interact with EGFP-tagged WT RNF213 (**Fig. 3a,b**). The AAA+ ATPase mutant alters the glutamic acid residue in the first Walker B motif (E2488), preventing ATP hydrolysis, which is required for homo-hexamer disassembly. As expected, FL-RNF213^WT^ co-immunoprecipitated with EGFP-RNF213. Mutations in the AAA+ ATPase or RING domain did not alter binding to EGFP-RNF213; likewise, RNF213^R4810K^ retained oligomerization activity (**Fig. 3a,b**). Because FL-RNF213^I3999A^ and FL-RNF213-WT associated comparably with EGFP-RNF213, E3-ligase activity is not required for, nor does it regulate, oligomerization.

**Figure 3:**
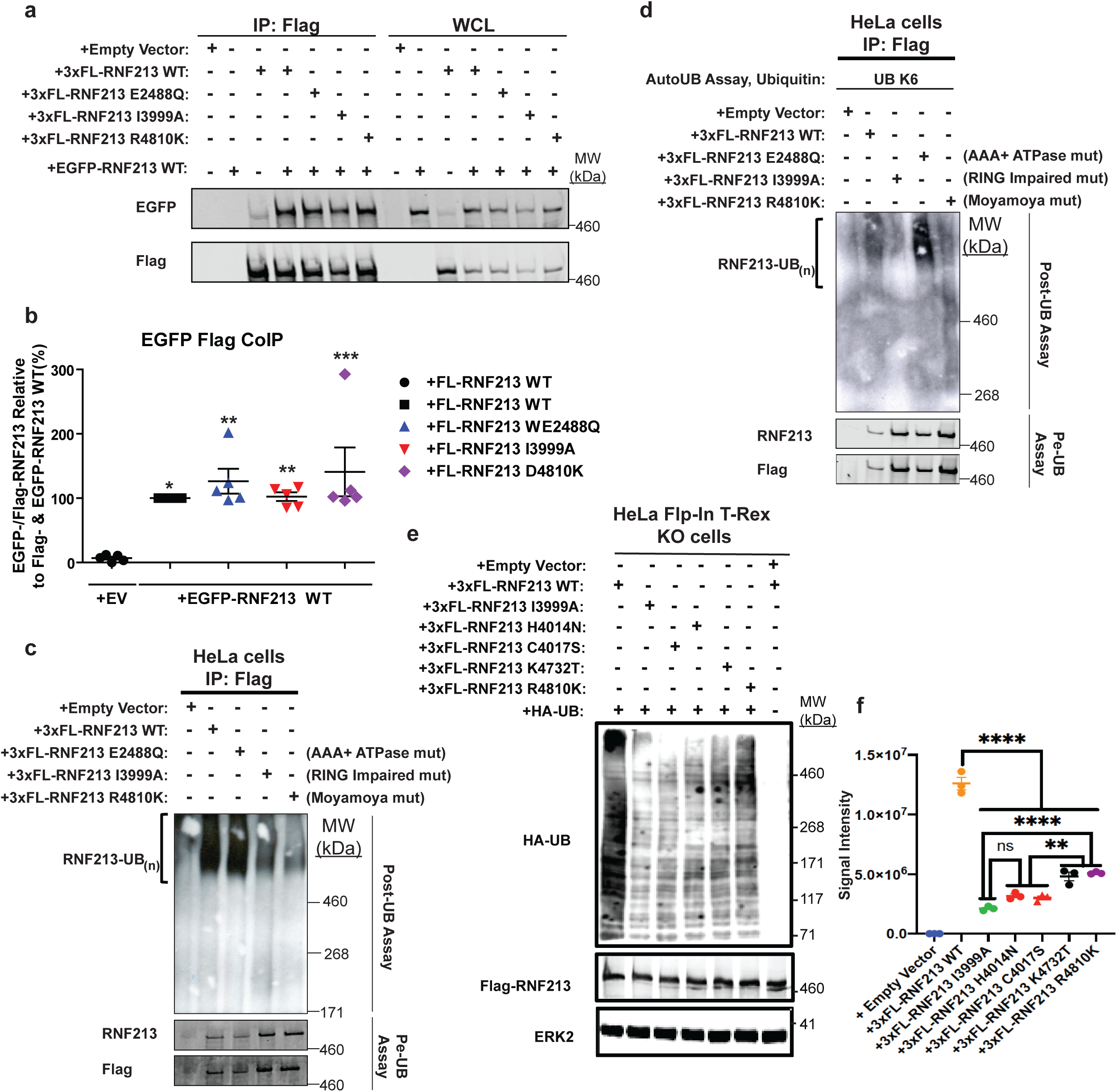
MMD-associated RNF213^R4810K^ has decreased E3-ligase activity. **a**, RNF213 mutants interact with RNF213^WT^. Lysates from 293T cells co-transfected with wild-type (WT) EGFP-RNF213 and 3xFL-RNF213-WT, -E2488Q, -I3999A or -R4810K were subjected to anti-Flag immunoprecipitation followed by anti-GFP immunoblotting. Expression of each RNF213 protein in whole cell lysates (WCL) is shown at right. **b**, Quantification of levels of EGFP-RNF213 WT co-immunoprecipitated with 3xFL-RNF213^WT^ and RNF213 mutants in **a. (***n=*5 biologically independent samples). Graph indicates mean percentage ± S.E.M. Statistical significance was evaluated by one-way ANOVA, followed by Bonferroni post-hoc test. **c**, *in vitro* auto-ubiquitylation assays of anti-Flag immunoprecipitates from lysates of transiently transfected HeLa cells expressing 3xFL-R*NF213*^*WT*^ or RNF213 mutants. **d**, Auto-ubiquitylation assays as performed in **c**, but in the presence of K6 ubiquitin. **e**, HeLa Flp-In T-Rex KO cells expressing 3xFL-RNF213^WT^ or the indicated RNF213 variants upon doxycycline induction were transfected with HA-UB expression construct. A portion of the cells were subjected to MG132 and Chloroquine for the last 3 hours prior to harvesting. Cells were harvested 36 hours post-transfection, and immunoblotting was performed for the indicated proteins. **f**, Quantification of immunoblots from **e** (n=3 biologically independent samples), which include the effects of RNF213 variants encoded by MMD SNPs with high penetrance, H4014N (red circles), low penetrance (purple circle) and undetermined penetrance (black circle), compared to a mutant in an essential RING domain cysteine, -C4017S(red triangle). Graph indicates integrated signal intensity ± S.E.M. Statistical significance was evaluated by one-way ANOVA, followed by Tukey test. Immunoblots (**c-e**) are representative images of at least three independent experiments.

Next, we performed *in vitro* auto-ubiquitylation assays using FL-RNF213-WT and FL-RNF213 variants purified from transiently transfected HeLa cells (**Fig. 3c**). As expected, auto-ubiquitylation was observed with FL-RNF213-WT, but this activity was decreased substantially in the RING-impaired mutant, FL-RNF213^I3999A^. Residual activity presumably reflects co-precipitating endogenous RNF213. The AAA+ ATPase mutation, *RNF213*^*E2488Q*^, did not affect RNF213 auto-ubiquitylation, indicating that ATP hydrolysis is not required for E3 ligase activity. By contrast, the MMD-associated RNF213^R4810K^ had auto-ubiquitylation activity, comparable to, if not lower than, RNF213^I3999A^ (**Fig. 3c**). We also asked whether full-length RNF213, like the RNF213 RING domain (**Fig. 1e**), can catalyze K6-Ubiquitin linkages. Indeed, using UBE2D2, FL-RNF213 WT and the AAA+ ATPase mutant (E2488Q), but not the RING-impaired (I3999A) or MMD-associated (R4810K) mutant, auto-ubiquitylated in the presence of K6-only UB molecules (**Fig. 3d**). Previously, it was reported that transiently expressed RNF213^R4810K^, but not a RING-deleted mutant, was ubiquitylated to levels similar to RNF213^WT^ in 293T cells, which have low levels of RNF213 expression (Liu et al., 2011). However, our experiments show that when RNF213^R4810K^ is purified from cells, it has decreased auto-ubiquitylation activity. This apparent discrepancy suggests that, in cells, (an)other E3-ligase(s) can ubiquitylate RNF213 on the RING domain. We also find that AAA+ ATPase hydrolysis by the first Walker B motif is dispensable for RNF213 E3-ligase activity, implying that the R4810K variant impairs auto-ubiquitylation through a mechanism independent on its effects on the AAA+ ATPase domain. Different *RNF213* SNPs are associated with variable penetrance of MMD (Cecchi et al., 2014; Guey et al., 2017; Kamada et al., 2011; Liu et al., 2011; Sugihara et al., 2019). To explore the role of RNF213 E3 ligase activity on MMD, we assessed the E3 ligase activity associated with SNPs of different penetrance **(Fig. 3e,f)**. Notably, all MMD SNPs encoded proteins with lower E3 activity than WT RNF213. Furthermore, RNF213^H4014N^ (higher penetrance, red circles) had lower activity compared with RNF213^R4810K^ (lower penetrance, purple circles). It has been reported that MMD SNPs in the RING domain of RNF213 have higher disease penetrance (Cecchi et al., 2014; Guey et al., 2017). Intriguingly, RNF213^H4014N^ and RNF213^C4017S^ affect RING domain residues in RNF213 and have highly impaired E3 ligase activity, compared with RNF213^R4810K^. Taken together, we conclude that MMD-associated *RNF213* SNPs impair E3 ligase activity. Furthermore, disease penetrance might depend upon the extent to which RNF213 E3 ligase activity is attenuated by a given MMD SNP.

### RNF213^R4810K^ acts as a dominant negative mutant in cells

RNF213 has global effects on the ubiquitylome of HER2+ breast cancer cells (Banh et al., 2016) and HeLa Flp-In T-Rex cells (**Fig 1h**). To assess the effect of MMD-associated RNF213 on cellular ubiquitylation, HA-tagged ubiquitin (HA-UB) was co-transfected with *RNF213* or *RNF213* mutants into 293T and HeLa cells. A portion of each transfected cell population was treated with proteasomal (MG132) and lysosomal (chloroquine) inhibitors to block UB-mediated degradation. Consistent with our previous findings, over-expression of WT RNF213 resulted in an increased number of ubiquitylated proteins relative to control, vector-transfected cells (**Fig. 4a-d**). Proteosomal and lysosomal inhibition resulted in a further increase in high molecular weight (>150kDa) ubiquitylated species. These global changes in ubiquitylation were dependent on RNF213 E3 activity, as shown by cells transfected with the RING mutant, *RNF213*^*I3999A*^. Instead, RNF213^I3999A^ expression actually *decreased* overall ubiquitylation compared with vector-transfected cells. Expression of the AAA-ATPase mutant (E2488Q) increased overall ubiquitylation to an extent similar to that of WT RNF213, indicating that ATPase activity is dispensable for RNF213 E3 action in cells, as well as *in vitro*.

**Figure 4:**
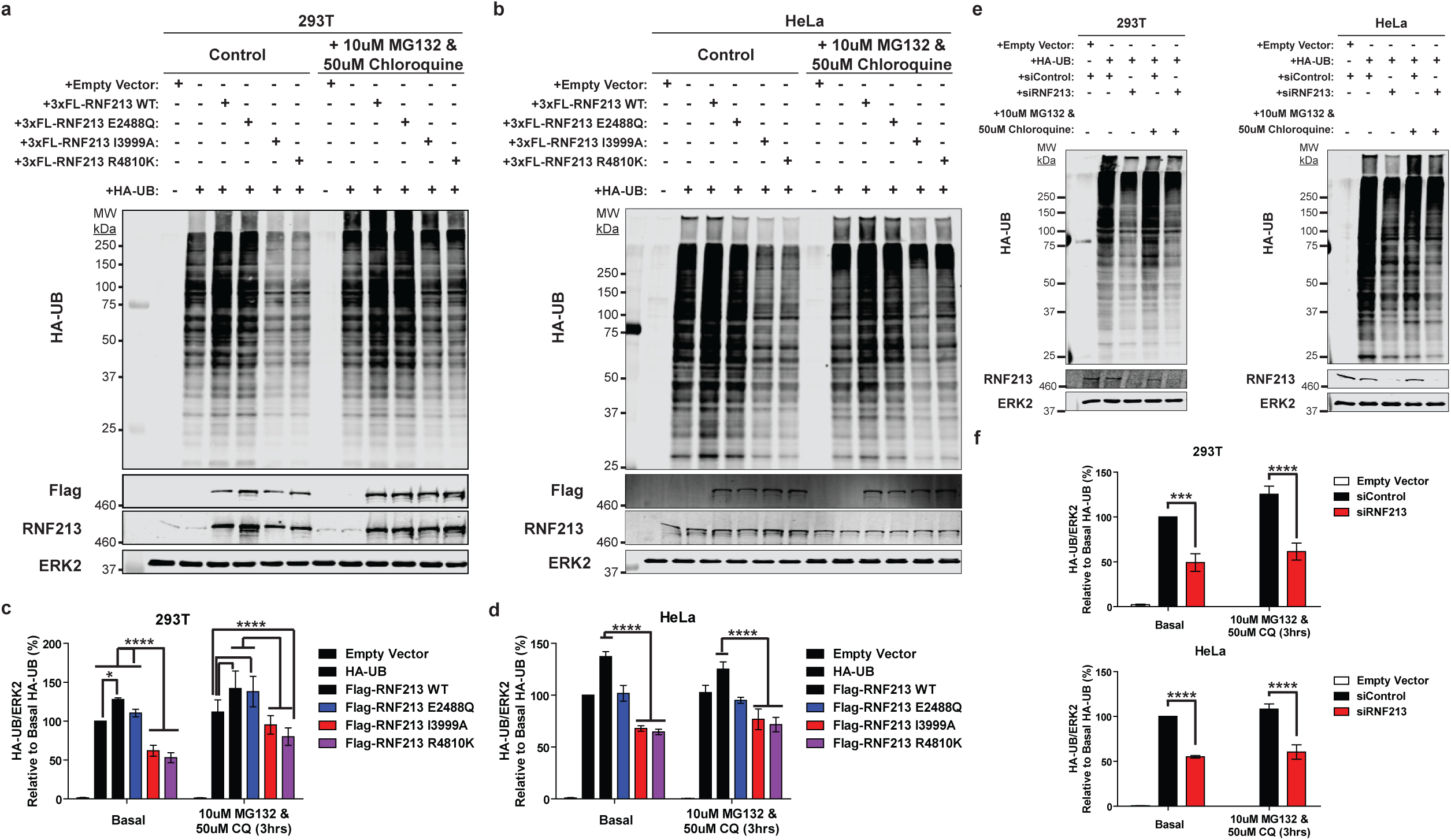
E3 ligase-impaired RNF213 acts as a dominant-negative mutant that affects global ubiquitylation. Immunoblots for HA-UB from 293T (**a**) and HeLa (**b**) cells transiently co-transfected with HA-UB and 3xFlag-RNF213^WT^ or RNF213 mutant expression constructs. A portion of each transfected cell population was treated with the indicated concentrations of proteasomal (MG132) and lysosomal (chloroquine) inhibitors for 3hrs. **c** and **d**, quantification of immunoblots from **a** and **b**, respectively (*n=*5 and 6 biologically independent samples for HeLa and 293T cells, respectively). **e**, 293T and HeLa cells were transfected with Control or *RNF213* siRNAs, followed by an HA-UB expression construct, and treated with the indicated proteasome and lysosome inhibitors for 3 hrs. Lysates were subjected to anti-HA immunoblotting, and results are quantified in **f (***n=*4 biologically independent samples). ERK2 served as a loading control. Graphs indicate mean percentage of cells ± S.E.M. Statistical significance was evaluated by two-way ANOVA, followed by Bonferroni post-hoc test.

These results suggested that, consistent with its oligomeric structure, E3-defective RNF213 can act as a dominant negative mutant. Indeed, cells over-expressing *RNF213*^*R4810K*^, which also has decreased E3-ligase activity (**Fig. 3c, 4a-d**), exhibited a global decrease in ubiquitylation compared with controls and cells over-expressing RNF213^WT^ (**Fig. 4a-d**). We also assessed the effects of *RNF213* knockdown (*RNF213*-KD). Similar to the dominant negative effects of the RING-impaired and MMD-associated *RNF213* mutants, *RNF213-*depleted 293T and HeLa cells had a global decrease in ubiquitylation (**Fig. 4e, f**). These observations, together with the effects of *RNF213* perturbations in breast cancer cells (Banh et al., 2016), show that RNF213 plays a crucial role in regulating ubiquitylation in multiple cell types. In breast cancer cells, RNF213-dependent ubiquitylation involved multiple proteins involved in ubiquitin metabolism, including E1s, E2s, E3s and deubiquitylases (Banh et al., 2016). Most likely, the global increase in ubiquitylation caused by RNF213 over-expression in 293 or HeLa cells also reflects a combination of direct and indirect effects on the cellular ubiquitylation machinery.

## Discussion

RNF213 is particularly interesting because it contains AAA-ATPase and RING E3 ligase domains in the same molecule, is associated with MMD (Liu et al., 2011), and is required for the toxic effects of PTP1B deficiency/inhibition in the hypoxic HER2+ breast cancer cells (Banh et al., 2016). Previous studies showed that RNF213 oligomerizes via its ATPase domain (Morito et al., 2014) and has ubiquitin ligase activity (Banh et al., 2016; Kobayashi et al., 2015). ATPase activity also has been implicated in recruitment to, and stabilization of, lipid droplets (Sugihara et al., 2019). However, the type of UB linkages added by RNF213, the cooperating E2(s), and the role(s) of the AAA-ATPase and E3 domains in MMD pathogenesis have remained unknown or controversial. We find that, acting in concert with UBE2D2, RNF213 can catalyze K6-ubiquitylation. Furthermore, while RNF213 variants encoded by MMD alleles have apparently unaltered ATPase activity, their E3 ligase is impaired, and the extent of impairment is greater in more penetrant alleles. Consistent with its oligomeric structure, MMD alleles act as dominant negative alleles in cells, providing a potential explanation for the autosomal dominant inheritance pattern of this syndrome.

Little is known about the function of K6-UB linkages in cells. Only three other ubiquitin ligases in mammalian cells are known to mediate K6-ubiquitylation: PARKIN, HUWE1 and BRCA1:BARD1 heterodimer (Michel et al., 2017; Wu et al., 2008; Yau and Rape, 2016). K6-UB linkages contribute to PARKIN-dependent mitophagy (Yau and Rape, 2016). In recent work, HUWE1 was found to regulate K6-UB linkages on mitofusin-2 (Michel et al., 2017). BRCA1:BARD1, which assembles K6-linkages on itself and its substrates (nucleophosmin, CTIP (C-terminal binding protein 1) and RNA polymerase subunit RPB8), plays a key role in DNA replication and homologous recombination repair. K6-poly-ubiquitylated BRCA1 is recognized, recruited and stabilized at sites of DNA damage by RAP80 (receptor-associated protein 80). BRCA1 K6-linkages also recruit UBXN1 (UBX domain-containing protein 1), which binds to, and acts as a cofactor for, p97 VCP (valosin-containing protein) to negatively regulate BRCA1:BARD1 E3-ligase activity (Wu et al., 2008). Notably, p97 VCP is an AAA-ATPase (Meyer et al., 2012); in this context, ATPase and E3 ligase activity reside in a single polypeptide in RNF213, whereas these functions are divided among several species in the p97 VCP/BRCA1:BARD1 interaction. PARKIN HUWE1 and BRCA1:BARD1 also can promote other ubiquitin linkages to form heterotypic chains. Although our results reveal a role for UBE2D2 in the control of RNF213 activity, its specific, physiological role awaits the identification of direct RNF213 substrates. With the recent advances in the detection of K6-UB-linkages using affimers, it will be interesting to study RNF213-dependent K6-ubiquitylated substrates (Michel et al., 2017).

Ubiquitin conjugating enzymes play a key role in the ubiquitylation process. Humans have ∼ 40 ubiquitin conjugating enzymes that are involved in ubiquitinylation or ubiquitin-like modifications of proteins (Stewart et al., 2016). Within cells, E2s exist primarily as E2-UB conjugates and are thus, in principle, capable of directly ubiquitylating substrate proteins. However, reactivity is low in the absence of an appropriate E3. HECT domain-containing E3 ligases transfer UB via E3-Ub intermediates, rendering linkage specificity dependent predominantly on the specific E3. By contrast, RING E3s, such as RNF213, require E2-UB conjugates to facilitate UB transfer. Hence, the specific types of UB linkages catalyzed by RING E3s is determined by the associated E2. Our results suggest that UBE2D2 mediates K6-UB linkages via RNF213. Notably, a previous report suggested that BRCA1:BARD1 promotes K6-ubiquitylation via UBE2D3 (Nishikawa et al., 2004; Wu-Baer et al., 2003). We reported previously that that PTP1B-deficiency or –inhibition increases RNF213 E3 ligase activity (Banh et al., 2016). In that study, di-Gly antibody-based UB proteomics revealed increased K6/11, K29, and K33 ubiquitylation in PTP1B-deficient cells (Kim et al., 2011). Consistent with the *in vitro* results above, those increases were normalized by *RNF213* knockdown. Taken together, these results suggest that UBE2D family of ubiquitin-conjugating enzymes plays an important role in catalyzing K6-UB linkages. A recent study argued that RNF213, in concert with UBE2N/UBE2V1, generates K63 ubiquitin linkages (Takeda et al., 2020). Given that the linkage specificity of RING E3 ligases is determined primarily by the E2s with which they interact, RNF213 might generate different linkages with different E2s. Therefore, the physiological relevance of E2s in context with the E3 ligase activity is important. Our knockdown data indicate that, at physiological levels, UBE2D2 is a relevant E2 for RNF213 E3 ligase activity, at least in HeLa Flp-In Tex cells.

We also find that MMD-associated SNPs impair RNF213 E3 ligase activity, but not ATPase activity. Furthermore, our results suggest that the penetrance of different SNPs might reflect the degree to which then allele impairs E3 ligase activity. Studies of larger panels of RNF213 alleles will be required to solidify these findings. A recent study suggested that RNF213 localizes to and stabilizes lipid droplets (Sugihara et al., 2019). Furthermore, RNF213 also stabilizes the droplets by removing ATGL (a lipase). Intriguingly, ATPase and E3 ligase activity of RNF213 both were shown to be important for stable association with lipid droplets, although E3 activity was dispensable for ATGL expulsion. Moreover, MMD SNPs with high penetrance (C3997Y, H4014N, C4017S, and C4032R) had more impaired lipid droplet targeting/droplet stabilization than MMD SNPs with low penetrance (D4013N, R4810K). Our finding that MMD penetrance correlates with the degree of impairment of RNF213 E3 ligase activity comports with the notion that aberrant regulation of RNF213 association with lipid droplets could be central to MMD pathogenesis.

At first glance, the finding that RNF213^R4810K^ has impaired E3 ligase activity would seem inconsistent with the autosomal dominant inheritance of MMD. However, we find that this MMD variant, as well as an engineered RING domain mutant, has dominant negative effects. Like RNF213^WT^, RNF213^R4810K^ and RNF213^I3999A^ retain the ability to oligomerize (Morito et al., 2014), indicating that decreased auto-ubiquitylation activity cannot be explained by inability to form a homo-hexamers. Similarly, E3 activity also does not depend on RNF213 AAA+ ATPase activity. Taken together, our findings provide a biochemical explanation for the observed dominant negative effects of E3-defective RNF213 mutants in cells. As in other multimeric complexes, such as p97VCP and RVB1/2 (RuvB family ATP-dependent DNA helicase pontin), incorporation of one or a few defective subunits to the RNF213 holoenzyme probably disables the entire complex (Jha and Dutta, 2009; Tresse et al., 2010). Further work, including structural analysis, is needed to determine how a single MMD SNP, outside of the RING and AAA+ ATPase domains, impairs the E3 activity of the other complex members

The dominant negative effects of RNF213^R4810K^, the defective E3 ligase activity of other prevalent MMD SNPs, and the evidence that penetrance and degree of E3 ligase impairment might correlate, all indicate that defective E3 ligase activity probably is important in MMD pathogenesis. Consistent with this hypothesis, several other MMD-associated SNPs directly affect the RING domain, including *RNF213*^*C3997Y*^, which alters a key cysteine that positions the zinc atom in the RING and should abrogate E3 ligase activity. It is therefore surprising that MMD-like phenotypes are absent in *Rnf213-/-* or *Rnf213*^R4828K/+^ mice. Furthermore, approximately 2% of the Japanese population have *RNF213* SNPs, yet the incidence of MMD in Japan is as low as 0.53 per 100,000. Notably, incidence is higher in children and adolescents (5-14 yrs old) and is ∼2-fold more common in females than in males. MMD also is associated with anemia, autoimmunity, infection/inflammation, and radiation. Together, these findings predict that genetic or environmental modifier(s) are required to trigger MMD in patients bearing *RNF213* SNPs. It will be important to identify such modifiers and determine how they affect RNF213 E3 activity or its substrates. A clue might come from considering biological differences between mice and fish, as *Rnf213* morphants in zebrafish do show abnormal vascular growth (Liu et al., 2011).

## Acknowledgements

This work was funded by R01 CA49152 to B.G.N. Work done in the S.S.S. lab for this study was supported by a Genome Canada Disruptive Innovation in Genomics grant (OGI-119) and a CIHR operating grant (MOP#136956). W.Z. is the recipient of the Cancer Research Society/ BMO Bank of Montreal Scholarship for the Next Generation of Scientists. R.S.B. was supported by a Doctoral Fellowship Grant from the Canadian Breast Cancer Foundation. W.Z. was supported by a CIHR Post-doctoral Fellowship Grant.

## Contributions

A.B., R.S.B. and W.Z. designed and performed most of the experiments. S.S.S. provided conceptual advice on E3 ligases. B.G.N. and S.S.S. conceived and supervised the project and helped to interpret the data. A.B., R.S.B. and B.G.N. wrote the manuscript. All authors critically analyzed and approved the data and edited the manuscript.

## Materials and Methods

### Cell lines and culture conditions

HeLa Flp-In T-Rex, HeLa and 293T cells were grown in DMEM with 10% FBS (Invitrogen), 100 U/mL penicillin, and 100 mg/mL streptomycin at 37°C. HeLa Flp-In T-Rex were purchased from Thermo Fisher Scientific. HeLa and 293T cell lines were obtained from the ATCC, authenticated by STR testing, and assessed monthly for the absence of mycoplasma contamination (MycoAlert, Lonza). Transfections were carried out by using Fugene 6 (Promega), according to the manufacturer’s instructions. In transient transfection experiments, the amount of plasmid DNA was kept constant between conditions in every experiment by adding appropriate amounts of empty vector. Control siRNAs (Cat. No. D-001810-10-05) or targeting *RNF213* (Cat. No. L-023324-00-0005) or *UBE2D2* (Cat. No. L-010383-00) (Dharmacon) were introduced into cells using Lipofectamine RNAiMAX (Thermo Fisher Scientific.), as per the manufacturer’s instructions.

An *RNF213*-knockout HeLa Flp-In T-Rex cell line was generated using CRISPR/Cas9 technology (Ran et al., 2013). Briefly, an sgRNA targeting the third exon of *RNF213* (ACAATGGCGTCGGCCTCGGA) was designed by using http://crispr.mit.edu and cloned into the BbsI site of pSpCas9 (BB)–2A–Puro (PX459, Addgene). HeLa cells were transiently transfected with the PX459–sgRNF213-exon3 vector using FuGENE 6 Transfection Reagent (Promega). Forty-eight hours post-transfection, cells were diluted and plated in 96 well plate at a density of 1 cell/well. Homozygous RNF213 knockout clones were identified by DNA sequencing and confirmed by immunoblotting. For generating isogenic HeLa Flp-In T-Rex knockout cells expressing WT RNF213 and RNF213 variants upon doxycline induction, pcDNA5/FRT/TO-based constructs expressing 3x-Flag tagged RNF213 or RNF variants were co-transfected with pOG44, which directs constitutive expression of Flp recombinase. Cells with successful integration of 3x-Flag-RNF213 or its variants were selected by adding hygromycin B (200 µg/mL) at 48 hours post-transfection. RNF213 expression was induced by adding 1 µg/mL doxycycline.

### Immunoblotting and immunoprecipitation

Cells were lysed in lysis buffer [50mM Tris·HCl (pH 8), 150 mM NaCl, 2 mM EDTA, 10mM Na4P2O7, 100 mM NaF, 2 mM Na3VO4, 1% (vol/vol) NP-40, 40 µg/mL phenylmethyl sulfonyl fluoride, 2 µg/mL antipain, 2 µg/mL pepstatin A, 20 µg/mL leupeptin, and 20 µg/mL aprotinin]. For whole cell lysate immunoblots, equal amounts of protein per sample were subjected to SDS-PAGE and transferred to a PVDF membrane (Millipore). For immunoprecipitations, lysates were incubated with appropriate antibody-agarose beads, as indicated, for 4 hours on a rotator-mixer at 4°C. Beads were washed 5 times in lysis buffer, and then analyzed by immunoblotting. To detect RNF213, lysates and immunoprecipitates were resolved by modified SDS-PAGE, using 3-8% Tris-acetate gels or 5% Tris-glycine gels. Gels were transferred in 1X transfer buffer, 2% methanol for 16 hours at 25 Volts. The following antibodies were used: RNF213 (Millipore, Cat. No. ABC1391), HA (Cell Signaling Technology, Cat. No. 3724), GST (Santa Cruz, Cat. No. sc-138), Flag (Sigma, Cat. No. F1804) ERK2 (Santa Cruz, Cat. No. sc-1647) GFP (Cell Signaling Technology, Cat. No. 2956). Blots were quantified by acquisition software, Image Studio Lite, using an Odyssey Infrared imaging system (Li-Cor Biosciences). Effects on overall ubiquitylation in **Fig. 3e,f** and **Fig. 4a-f** were assessed by integrating the intensity of the HA-UB signal in each lane.

### *In vitro* ubiquitination assays

Recombinant RNF213 RING domain, fused to GST (GST-RNF213^RING^) protein was purified from *E. coli* strain BL21. Assays of GST-RING^RNF213^ activity were performed by using the E2Select Ubiquitin Conjugation Kit (Boston Biochem, Cat. No. K-982), according to the manufacturer’s instructions. Briefly, recombinant RNF213 was incubated in a reaction buffer containing E1, E2 (E2Select Ubiquitin Conjugation Kit), and ubiquitin, at 37’C for 1 hour. Reactions were terminated by adding SDS-PAGE sample buffer, and ubiquitin incorporation was analyzed by immunoblotting, as described above.

### ATPase Assays

ATPase assays using full length RNF213 were carried out using the ATPase Activity Assay Kit (Biovision, Cat. No. K417-100), as per the manufacturer’s instructions. Briefly, purified RNF213 (3x-Flag-RNF213^WT^, 3x-Flag-RNF213^E2488Q,E2845Q^, 3x-Flag-RNF213^I3999A^, 3x-Flag-RNF213^C3997Y^, 3x-Flag-RNF213^D4013N^, 3x-Flag-RNF213^H4014N^, or 3x-Flag-RNF213^R4810K^, as indicated) was incubated in assay buffer containing ATPase substrate, developed at 25°C for 15-30 minutes, and A_650_ was measured using FlexStation 3 Multi-Mode Microplate Reader. For full-length RNF213 purification, HeLa Flp-In T-Rex KO expressing 3x-Flag tagged wild-type RNF213 or RNF213 variants were induced with doxycycline (1µg/mL) and lysed in ice cold ATPase Assay Buffer provided with the ATPase assay kit. Lysates were subjected to immunoprecipitations with anti-FLAG M2 magnetic beads (Sigma, Cat. No. M8823) for 3 hours on a rotator-mixer at 4°C. Beads were collected at bottom of the tube using a magnetic stand, supernatants were discarded, and immunoprecipitates were washed 5 times in lysis buffer followed by elution with 50µL of 100ng/µL 3x-Flag peptide.

### Ubiquitylome analysis

Where indicated, cells were transfected with empty pRK5 vector or pRK5 expressing HA-UB (HA-tagged ubiquitin) by using Fugene 6 (Promega), as per the manufacturer’s instructions. In some experiments, a portion of the transfected cells was treated with 10 µM MG132 (Tocris) and 50 µM chloroquine (Tocris). Cells were lysed in lysis buffer [50mM Tris·HCl (pH 8), 150 mM NaCl, 2 mM EDTA, 10mM Na4P2O7, 100 mM NaF, 2 mM Na3VO4, 1% (vol/vol) NP-40, 40 µg/mL phenylmethyl sulfonyl fluoride, 2 µg/mL antipain, 2 µg/mL pepstatin A, 20 µg/mL leupeptin, and 20 µg/mL aprotinin] along with deubiquitinase inhibitor (+10mM IAM and NEM), and lysates were analyzed by immunoblotting as described above.

### Quantitative real-time PCR

Total RNA was extracted using miRNeasy kit (Qiagen, Cat. No. 217004). cDNA synthesis was carried out using the RT2 first stand kit (Qiagen) cDNA synthesis kit following the manufacturer’s protocol. qPCR then was carried out using Maxima SYBR Green Supermix (Life Technologies) in an Applied Biosystems StepOne Real-time PCR machine. 18S rRNA was used as the internal control for all samples. For human *UBE2D2*, following primers were used: *GGTCACAGTGGTCTCCAGCACTAA* and *ACTTCTGAGTCCATTCCCGAGCT.*

### Statistical Analysis

Sample sizes and statistical tests for each experiment are mentioned in the figure legends. All analyses and graphs were generated using GraphPad Prism 8.

