## Supplementary figures and images for "Moyamoya Disease-Associated *RNF213* Alleles Encode Dominant Negative Alleles That Globally Impair Ubiquitylation"

### Supplementry Figure 1

**Supplementry Figure 1**

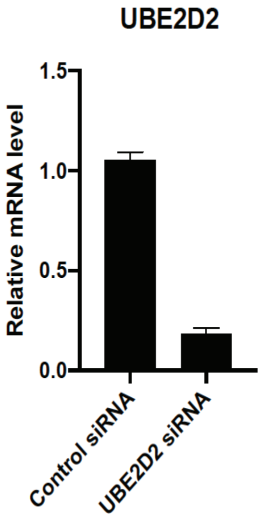
